# Efficient approaches for large scale GWAS studies with genotype uncertainty

**DOI:** 10.1101/786384

**Authors:** Emil Jørsboe, Anders Albrechtsen

## Abstract

1

**Introduction:** Association studies using genetic data from SNP-chip based imputation or low depth sequencing data provide a cost efficient design for large scale studies. However, these approaches provide genetic data with uncertainty of the observed genotypes. Here we explore association methods that can be applied to data where the genotype is not directly observed. We investigate how using different priors when estimating genotype probabilities affects the association results in different scenarios such as studies with population structure and varying depth sequencing data. We also suggest a method (`ANGSD-asso`) that is computational feasible for analysing large scale low depth sequencing data sets, such as can be generated by the non-invasive prenatal testing (NIPT) with low-pass sequencing.

**Methods:** `ANGSD-asso`’s EM model works by modelling the unobserved genotype as a latent variable in a generalised linear model framework. The software is implemented in C/C++ and can be run multi-threaded enabling the analysis of big data sets. `ANGSD-asso` is based on genotype probabilities, they can be estimated in various ways, such as using the sample allele frequency as a prior, using the individual allele frequencies as a prior or using haplotype frequencies from haplotype imputation. Using simulations of sequencing data we explore how genotype probability based method compares to using genetic dosages in large association studies with genotype uncertainty.

**Results & Discussion:** Our simulations show that in a structured population using the individual allele frequency prior has better power than the sample allele frequency. If there is a correlation between genotype uncertainty and phenotype, then the individual allele frequency prior also helps control the false positive rate. In the absence of population structure the sample allele frequency prior and the individual allele frequency prior perform similarly. In scenarios with sequencing depth and phenotype correlation `ANGSD-asso`’s EM model has better statistical power and less bias compared to using dosages. Lastly when adding additional covariates to the linear model `ANGSD-asso`’s EM model has more statistical power and provides less biased effect sizes than other methods that accommodate genotype uncertainly, while also being much faster. This makes it possible to properly account for genotype uncertainty in large scale association studies.

## 2 Introduction

Genome-wide association studies (GWAS) have classically been done to study genotype phenotype associations. However, a slightly different design is using low depth next-generation sequencing (NGS) data, because in such cases the genotype cannot be inferred accurately. An example of this study design is the non-invasive prenatal test (NIPT) for fetal trisomy by low-pass sequencing that provides a cost efficient design, where the number of individuals studied can be maximised, as each individual will be a lot cheaper to sequence when only low depth NGS sequencing is needed. The statistical power to detect associations increases with the number of individuals and therefore this design provides good statistical power to detect associations [Pasaniuc et al., 2012]. Another approach that is commonly used in GWAS is doing haplotype imputation to infer missing genotypes, this also generates genetic data with uncertainty on the observed genotype.

A recent successful GWAS [Liu et al., 2018] with low depth NGS data has shown the viability of this approach. In Liu et al. [2018] around 140,000 individuals had a NIPT for fetal trisomy by low-pass sequencing to an average sequencing depth of 0.1*X* and subsequently haplotype imputation was performed on the low depth NGS data. For the association testing a score test approach using a linear model framework [Skotte et al., 2012] implemented in ANGSD [Korneliussen et al., 2014] was used, as this method takes genotype uncertainty into account. Despite the low sequencing depth several novel associations were discovered. This provides an example of a study where using methods that account for the genotype uncertainty in low depth NGS data, provide good statistical power for detecting associations, despite the modest amount of data.

Using methods that take genotype uncertainty into account have advantages compared to calling genotypes for low depth NGS data, and then doing association analysis with those, as shown in Skotte et al. [2012]. In that method, implementing a score test, the coefficients are not estimated under the alternative hypothesis making the method computationally very fast, however this means the effect size of the genotype is not estimated. In this paper we will introduce `ANGSD-asso`, it works in a generalised linear model framework that also estimates the effect size of the unobserved genotype, and that in practice can be run almost as fast as the score test. `ANGSD-asso`’s EM model uses a maximum likelihood approach, more specifically we will make use of the EM algorithm to maximise the likelihood, treating the unobserved genotype *G* as a latent variable. Using a generalised linear model framework enables us to include covariates thereby adjusting for possible confounders, such as population structure. We have implemented an EM algorithm that converges fast and that can be run multi-threaded, making the analysis of large data sets possible. We have also implemented a hybrid model in `ANGSD-asso` combining the speed of the score test with the desired properties of the EM algorithm based approach. Using the EM algorithm for doing maximum likelihood estimation in a generalised linear model framework using genotype probabilities has been implemented in SNPTEST [Marchini et al., 2007]. We have designed a faster implementation that allows for the association analysis of large scale NGS data sets.

A common practice for doing association analysis with genotype data with uncertainty is using genetic dosages, which is the expected genotype calculated from the genotype probabilities. Dosages are easy to implement into most existing methods as the genotype can be directly replaced by the dosage. However dosages do not convey the uncertainty on the genotype as fully as genotype likelihoods or genotype probabilities. In Zheng et al. [2011] they show a gain in power when using genotype probability based methods compared to dosages, but only for small studies with variants with large effect sizes. However, they did not look at how a correlation between the sequencing depth and the phenotype might affect this. This could happen in a case-control study, where a systematic bias in the sequencing depth could be generated if cases and controls were sequenced at different places, or if the total data set is merged from other smaller heterogeneous data sets. We will investigate this sequencing depth bias through simulations of a large scale association study with a sequencing depth bias. We will evaluate the performance of our genotype probability based method compared to using dosages. In order to be able to asses how a sequencing depth bias might impact the analyses, we focus on simulating genetic data as low depth NGS data. However, we show that a scenario similar to this could also happen with imputation.

`ANGSD-asso`, and the score test and SNPTEST all work on genotype probabilities (also known as the “posterior probability”). The genotype probabilities can be calculated from the genotype likelihoods, which in turn can be calculated from the NGS data [Nielsen et al., 2011]. For more on genotype likelihoods and NGS data see Supplementary Section 4.2.

Another aspect we will explore is how to estimate genotype probabilities taking population structure into account, when doing association studies with low depth sequencing data. Population structure is a common confounder in association studies if not addressed properly [Marchini et al., 2004, Freedman et al., 2004, Cardon and Palmer, 2003]. In low depth sequencing data using a prior for calculating genotype probabilities based on the allele frequency estimated from the sequencing data (we will refer to this as the “sample allele frequency”) is often used, however this assumes a homogeneous population without structure. We therefore propose a novel method for dealing with structured populations, when doing association studies with low depth sequencing data. We will do this by using an individual allele frequency prior for when estimating the genotype probabilities. The individual allele frequency is the weighted average population frequency, the admixture proportions for each individuals are the weights. The individual allele frequency for a site has to be calculated for each individual, as their admixture proportions might differ. This takes both the frequency of the variant and the ancestry of every individual into account. We therefore want to investigate how this novel approach compares to using the sample allele frequency prior in different scenarios. We will look at this both with regards to statistical power and with regards to the false positive rate, and if there is correlation between the phenotype and the sequencing depth. As this scenario is most useful when working with data in non-model organisms, where good haplotype imputation cannot be done. We will do our simulations with low depth NGS data as our genetic data with uncertainty.

## 3 Methods

NGS produces short reads that are mapped to a reference genome. From the aligned reads the probability of observing these reads given a certain genotype can be inferred this is known as the genotype likelihood [Nielsen et al., 2011], for more on the genotype likelihood and how to calculate it see the Supplementary Section 4.2. The genotype likelihood can be converted into the probability of the genotype given the data, this is referred to as the genotype probability. For an overview of the relationship between the different kinds of genetic data, and how they can be processed and analysed in association see Figure 1.

**Figure 1:**
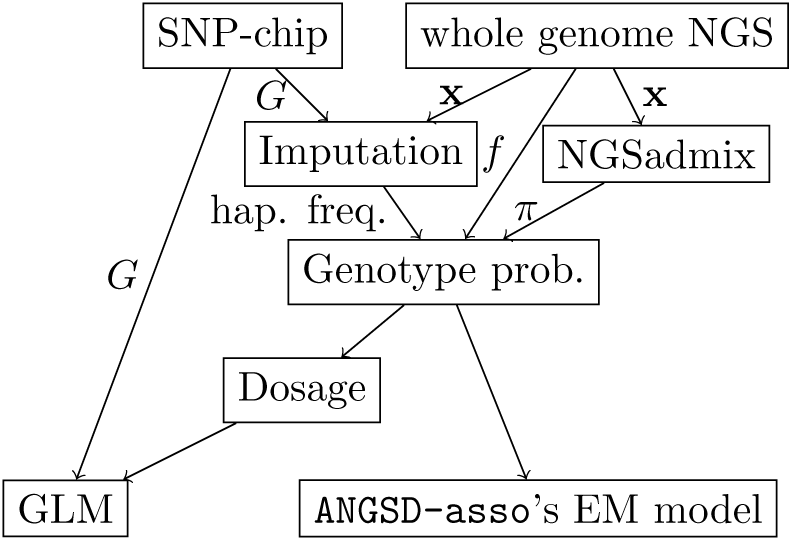
Schematic of workflow for doing association studies with genetic data. **x** is the sequence data that can be converted to genotype likelihoods, *G* is the genotypes. *π* is the individual allele frequencies and *f* is the sample allele frequency, these can both be used as priors when estimating the genotype probabilities. Data either gets generated using SNP-chips or doing whole genome NGS. The NGS data can be converted into genotype probabilities assuming no population structure using the sample allele frequency, or assuming population structure and then using PCA to generate genotype probabilities. SNP-chip genotypes can be analysed directly. Both kinds of data can be imputed using haplotype frequencies for generating genotype probabilities. The genotype probabilities can be analysed with `ANGSD-asso`’s EM model or converted to dosages and then be analysed with a generalised linear model (GLM).

### 3.1 ANGSD-asso’s EM model

We model the data using a maximum likelihood approach in a generalised linear model framework. This enables us to test for an association without observing the genotype *G* directly. Rather we observe our NGS data (**x**), from this we can infer *p*(*G*| **x**) or rather the probability of the genotype given the observed data (reads), this is also referred to as the genotype probability. We write the likelihood for our phenotype data (**y**) given our sequencing data (**x**) and covariates (*Z*)

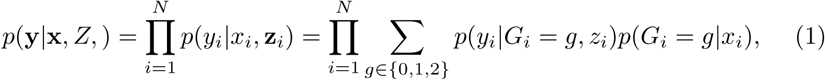

where we use the law of total probabilities to introduce the latent variable *G. N* is the number of individuals, **y** = (*y*_1_, *y*_2_, …, *y*_*N*_) is a vector of our observed phenotype for each individual, **x** = (*x*_1_, *x*_2_, …, *x*_*N*_) is a vector of sequencing data for each individual and *Z* = (**z**_**1**_, **z**_**2**_, …, **z**_**N**_) is a *N ×c* matrix with the covariates. We see that the trait *y*_*i*_ is conditionally independent of the sequencing data given the genotype (meaning *p*(*y*_*i*_ |*G*_*i*_ = *g, x*_*i*_, *z*_*i*_) = *p*(*y*_*i*_ |*G*_*i*_ = *g, z*_*i*_)). We can calculate the genotype probability only making use of the sequence data, for example by using the sample allele frequency as a prior, by assuming that the genotype is conditionally independent of the covariates, given the sequencing data and the frequency *f* (meaning *p*(*G*_*i*_ = *g*|*x*_*i*_, **z**_**i**_, *f*) = *p*(*G*_*i*_ = *g*|*x*_*i*_, *f*)), however for simplicity we omit *f* from the likelihood.

This allows us to write the likelihood, also introducing the parameters of our generalised linear model *θ* = (*α, β, γ*), again we assume that the genotype is conditionally independent of the covariates given the sequencing data

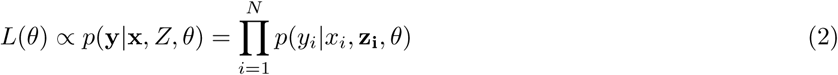

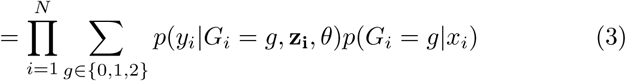

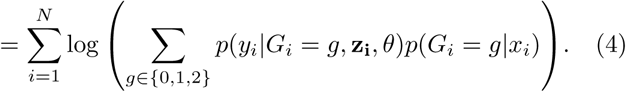

Assuming the term *p*(*y*_*i*_|*G*_*i*_ = *g*, **z**_**i**_, *θ*) follows a normal distribution, given the genotype *G* takes the value *g*, the covariates *Z* and the linear coefficients *θ*, the mean is given by

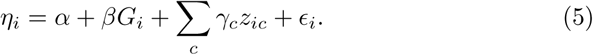

Equation 4 is the log-likelihood function that we want to maximise with regards to the parameters *θ*. For maximization we use the EM algorithm where our latent variable is the unobserved genotype *G*. This is done by weighted least squares regression, where the parameters *θ* are estimated. For the full derivations of this see the Supplementary Section 6 and 6.1. We have also implemented logistic and Poisson regression where we have introduced a link function for *η*_*i*_ for eq. 5 and changed the distribution for *p*(*y*_*i*_|*G*, **z**_**i**_, *θ*) accordingly. For more information on this see the Supplementary Section 5.

Furthermore standard errors on the estimated effect sizes are estimated using the observed Fisher information matrix as in Lake et al. [2003] and Skotte et al. [2019].

### 3.2 ANGSD-asso’s hybrid model - for fast computation

The score test as described in Skotte et al. [2012] only has to estimate the parameters of the null model, where uncertainty on the variables do not have to be taken into account. It is therefore faster than our approach, where we both have to estimate the null and the alternative model. The idea behind the hybrid model is combining the speed of the score test with the desirable properties of `ANGSD-asso`’s EM model, where estimates of the effect size and standard error can be obtained. It works by first running the score test, and then if the site has a P-value below a certain threshold, we additionally run the slower`ANGSD-asso`’s

EM model as well

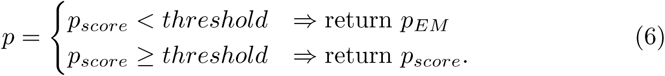

The threshold can be set by the user, the default value is 0.05.

### 3.3 ANGSD-asso’s dosage model

The expected genotype *E*[*G*|**x**] can easily be calculated from the genotype probabilities. This is an easy way to accommodate some of the genotype uncertainty and is therefore a method for trying to deal with genotype uncertainty in association studies

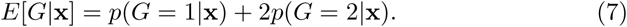

The genotype probability *p*(*G*|**x**) can be calculated using the genotype likelihood *p*(**x**|*G*) and the frequency *f* of the genetic variant, for calculating the prior *p*(*G*|*f*). This is done using Bayes’ formula

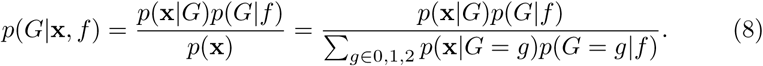

Here it is assumed that we have one homogeneous population where *f* describes the frequency of the genetic variant well across all individuals. Other priors can be used, for instance a prior based on the individual allele frequency (*π*) or haplotype frequencies. We do standard ordinary least squares using *E*[*G*|**x**] as our explanatory variable

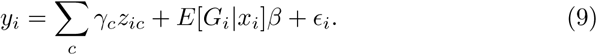

#### 3.3.1 Implementation

The three methods in `ANGSD-asso` for association analysis are implemented in the ANGSD framework [Korneliussen et al., 2014], allowing multi-threaded analysis. ANGSD can be downloaded from its github page: https://github.com/ANGSD/angsd The EM model is -doAsso 4, the hybrid model is -doAsso 5 and the dosage model is -doAsso 6. `ANGSD-asso` works on genotype probabilities in the BEA-GLE, BGEN and BCF/VCF file formats or directly from BAM files and the other formats allowed in ANGSD.

### 3.4 Individual allele frequency prior

When estimating the genotype probabilities for low depth sequencing data, it is important to have an accurate prior, when dealing with genotype data in a structured population. The sample frequency *f* of an allele might not describe the occurrence of an allele across individuals very well. This is due to the fact that the frequency of an allele might differ drastically between different ancestries. Therefore a prior based on the sample frequency will not work well in a structured population. If we have a discrete number of ancestral populations then by using a weighted average of the ancestral frequencies we can calculate the individual allele frequency (*π*_*ij*_), for individual *i* for site *j*, across *k* populations

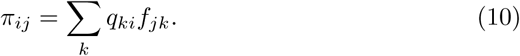

Where *f*_*jk*_ is the frequency of the *j*th site in population *k* and *q*_*ik*_ is the admixture proportion of population *k* for individual *i*. In order to estimate the individual allele frequencies we will have to first estimate the ancestral frequencies and the admixture proportions. For NGS data this can be done using NGSadmix [Skotte et al., 2013] and for genotypes this can be done using ADMIXTURE [Alexander et al., 2009]. We use the approach from NGSadmix when inferring population frequencies, in our simulations with low depth sequencing data in a structured population, assuming admixture proportions are known.

Another approach is [Hao et al., 2015] or PCAngsd [Meisner and Albrechtsen, 2018], where the population structure between individuals is modelled using principal components rather than a discrete number of ancestral populations. When the individual allele frequencies have been generated we can calculate more accurate genotype probabilities, this can be done using Bayes’ formula as laid out in eq. 8 (where we replace *f* by *π*). *p*(*G*) can be calculated using our individual allele frequency assuming Hardy-Weinberg proportions

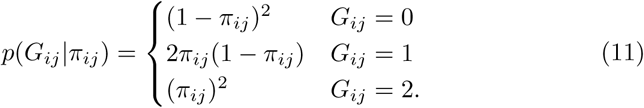

### 3.5 Simulated sequencing data

Sequencing data was simulated by first choosing an average depth for a group of individuals and then sampling the specific depth assuming a Poisson distribution. For simplicity we assumed a constant error rate of 1 %, furthermore we assumed that only two bases exist and sample the reads from these two alleles conditional on the simulated genotype and the error rate. For the run-times in Figure 5 and Supplementary Figure 12 and the effects of priming in Supplementary Figure 11, genetic data was simulated using frequencies from the Yoruba population from Lazaridis et al. [2014] where the curated Human Origins data set was selected.

We chose 6 different simulation scenarios, as summarised in Table 1. In scenario 1 we evaluate the false positive rate when there is sequencing depth and phenotype correlation, under our null hypothesis of no effect of the genotype. In scenario 2 we examine the statistical power when simulating under our alternative hypothesis with no sequencing depth and phenotype correlation. Scenario 3 is similar to scenario 1 and 4 is similar to 2, but with the addition of population structure. Scenario 5 and 6 are similar to scenario 2 and 4 respectively, but with correlation between sequencing depth and phenotype correlation. The sequencing depth and phenotype correlation was simulated using a logistic function, modelling the probability of being in the group with high average sequencing depth *p*(*D*_*i*_ = high).

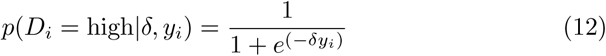

**Table 1:** Overview of simulations. We simulate under a standard additive model with a normally distributed phenotype with a mean given in the last column, and standard deviation 1. *g* is the genotype with effect *β* and *q* is the ancestry with effect *γ*. In Scenario 1 and 3 there is no effect of the genotype. In Scenario 3, 4 and 6 there is population structure, with two ancestral populations.

| Scenario | freq. | $N$ | population structure | seq. depth and pheno | simulated pheno mean |
| --- | --- | --- | --- | --- | --- |
| 1 | 0.45 | 1000 | no | correlated (eq. 12) | 0 |
| 2 | 0.45 | 1000 | no | not correlated | $\beta \cdot g$ |
| 3 | (0.9, 0.1) | 1000 | yes | correlated (eq. 12) | $q \cdot \gamma$ |
| 4 | (0.9, 0.1) | 1000 | yes | not correlated | $\beta \cdot g + q \cdot \gamma$ |
| 5 | 0.45 | 1000 | no | correlated (eq. 12) | $\beta \cdot g$ |
| 6 | (0.9, 0.1) | 1000 | yes | correlated (eq. 12) | $\beta \cdot g + q \cdot \gamma$ |

The higher (in absolute value) *δ* is, to a larger degree the phenotype will correlate with being in the group of high or low average sequencing depth, meaning the lowest values of the phenotype will have the lowest probability of being in the high depth group whereas the highest values of the phenotype will have the highest probability of being in the high depth group (if *δ* ¿ 0 and vice versa if *δ* ¡ 0).

From the simulation data we estimate frequencies from the genotype likelihoods. For the admixed individuals we assume that the admixture proportions are known, we estimate the population frequencies using the approach from Skotte et al. [2013]. The sequencing depth and phenotype correlation is simulated as described in eq. 12.

## 4 Results

In order to investigate what prior performs best for generating the genotype probabilities in different scenarios we simulated sequencing data with and without population structure and with and without sequencing depth and phenotype correlation. For each scenario we both applied a sample allele frequency prior and an individual allele frequency prior. We wanted to evaluate how both priors work with regards to false positive rates and statistical power to detect an association. Another aspect we wanted to investigate is statistical power in a large scale NGS association studies when using dosages versus when using our genotype probabilities based approach. We therefore simulated a large scale association study with low depth sequencing data. We also compared `ANGSD-asso`’s EM model with SNPTEST in terms of bias, statistical power and computational speed.

### 4.1 Evaluation of using different priors

#### 4.1.1 Using different priors in a homogeneous population

For scenario 1 with no population structure we explore the effect of a sequencing depth phenotype correlation. Supplementary Figure 1 shows an acceptable false positive rate in a population without structure for all approaches. Using an individual allele frequency prior and a sample allele frequency prior yield identical results. This is expected since these priors become identical in the absence of population structure. Supplementary Figure 2 shows that in scenario 2 there is no difference in statistical power between the two priors when there is no population structure.

#### 4.1.2 Using different priors in a structured population

For scenario 3 Figure 2 shows using the sample allele frequency prior gives us biased estimates of the effect size and leads to an increased false positive rate. The increased false positive rate is present even though we are adjusting for ancestry in the linear model, showing that this is not sufficient in this scenario. When using an individual allele frequency prior we do not get biased estimates and have a false positive rate that is the same as when using the true genotype.

**Figure 2:**
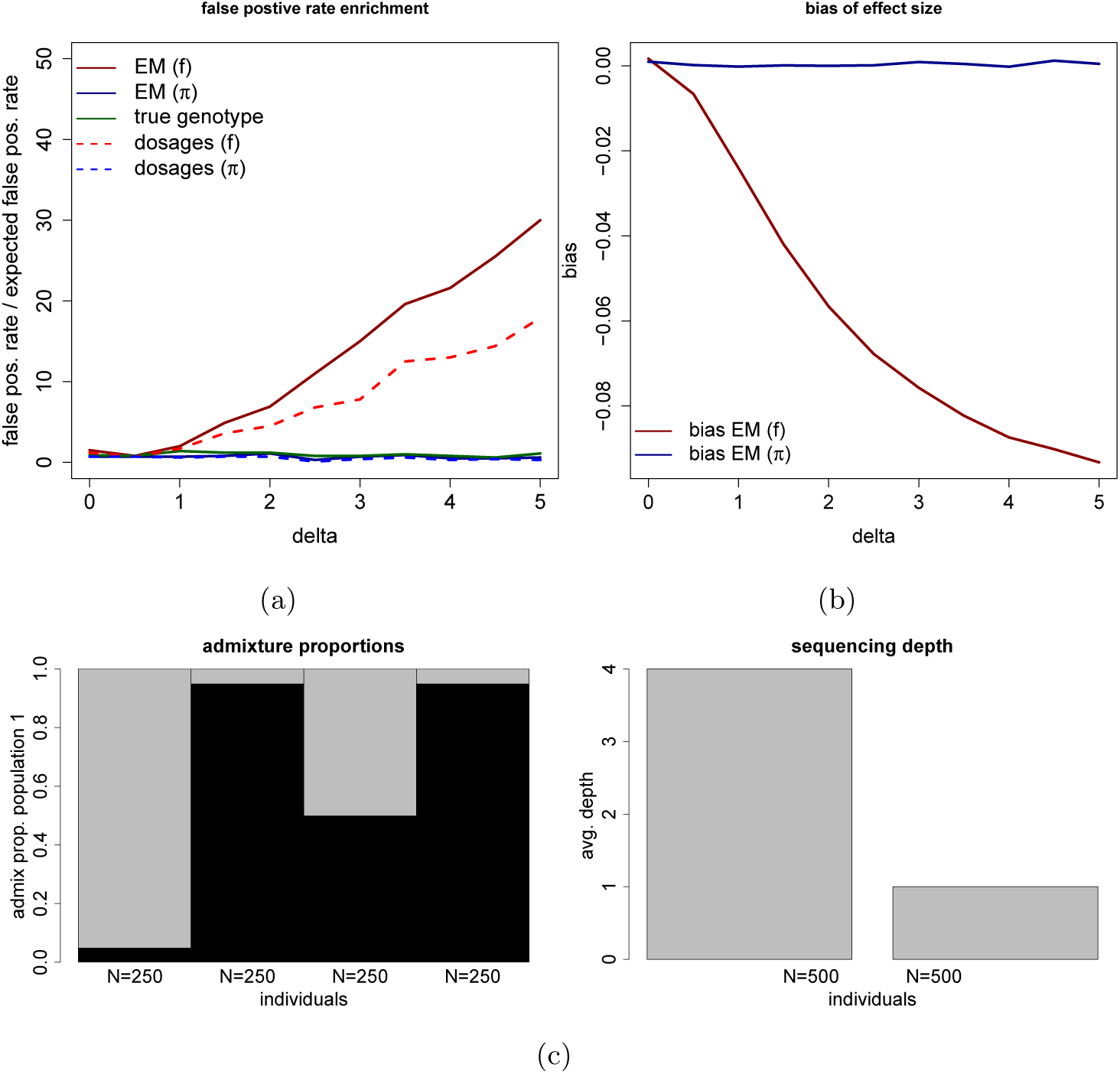
This data is simulated according to scenario 3 in Table 1 varying the sequencing depth and phenotype correlation (*δ*). We have a structured population with 1,000 individuals. There is an effect of ancestry of population 1 (*γ* = 1) which is adjusted for in the linear model. We use a significance threshold of 10^*–*3^. Each point is based on 10,000 simulations. **(a)**: We show the false positive rate divided by the expected false positive rate (10^*–*3^), using `ANGSD-asso`’s EM model and dosage model respectively with a sample frequency prior (f) and an individual allele frequency prior (*π*). **(b)** We show the bias of our estimated effect size. **(c)** The simulated admixture proportions and the mean sequencing depth for the simulated individuals.

For scenario 4 Figure 3 shows using our individual allele frequency approach leads to increased statistical power. For example for an effect size of *β* = 0.3 the power is 0.74 compared to 0.61, furthermore in this scenario we see that using dosages almost has similar statistical power to when using genotype probabilities. When using the sample allele frequency prior the effect sizes are underestimated. This is due to the fact that using the individual allele frequency better describes the expected genotype in a structured population.

**Figure 3:**
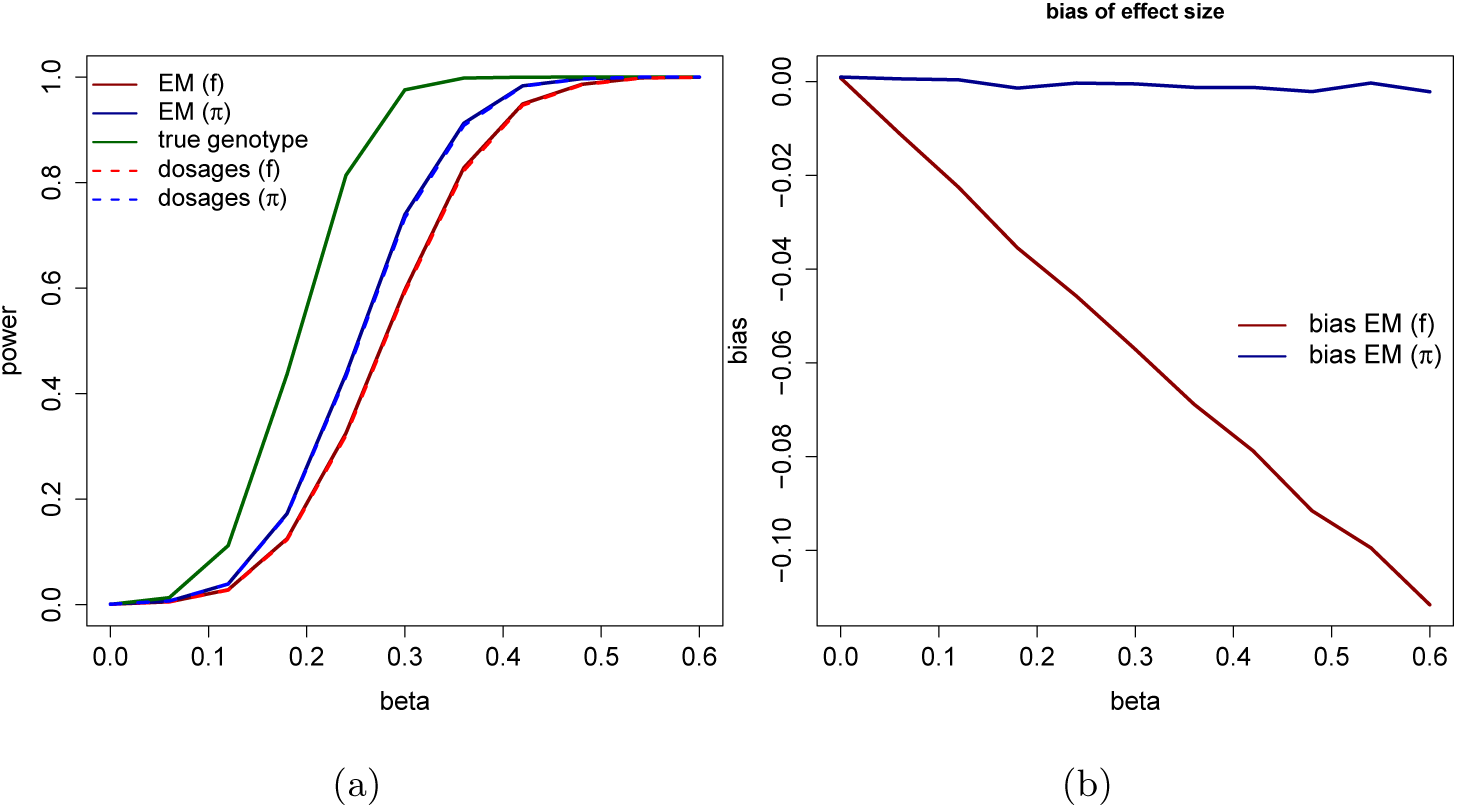
This data is simulated according to scenario 4 in Table 1 varying the effect size of the genotype (*β*). We have a structured population with the same admixture proportions and mean sequencing depth as in Figure 2 (c). There is an effect of ancestry of population 1 (*γ* = 1). We use a significance threshold of 10^*–*3^. The linear model is adjusted for ancestry. Each point is based on 10,000 simulations. **(a)**: We show the statistical power to detect a true association, using `ANGSD-asso`’s EM model and dosage model respectively with a sample frequency prior (f) and an individual allele frequency prior (*π*). **(b)**: We show the bias of our estimated effect size.

For scenario 5 in Table 1 that is like scenario 1, but with an effect of the genotype. Supplementary Figure 3b and 4b show that in this scenario there is slightly increased statistical power when using the full genotype probabilities compared to using dosages, and that our estimated effect size is less biased. For scenario 6 in Table 1 that is like scenario 3, but with an effect of the genotype. Supplementary Figure 5b and 6b show that in this scenario there is increased statistical power when using the full genotype probabilities compared to using dosages, and less bias of the estimated effect size.

### 4.2 Comparison with dosages in large scale studies

Genotype dosages or the expected genotype is often used in association studies, in order to be able to try and account for the uncertainty on the genotype. However dosages can be uninformative especially with low depth sequencing data. We did simulations in order to investigate the statistical power to detect an association, when we model the full genotype probabilities instead of using just the genotype dosage, using respectively `ANGSD-asso`’s EM and dosage model. We simulated a large case-control study with 100,000 individuals, to investigate a scenario with many individuals, as in the scenarios in Table 1 the number of individuals is much smaller. All individuals have low depth sequencing data, the cases and controls have been different average sequencing depths.

In Table 2 we show how using the full genotype probabilities have increased statistical power compared to when using the genotype dosages. We have more power for small effect sizes, where we have a true positive rate that is almost 0.1 higher. We calculated the info measure for our dosages in cases and controls respectively, to make it comparable with haplotype imputation. When genotypes are predicted with high certainty the info measure will be close to 1. We see that the info measure is lower in controls, where we have a lower average sequencing depth. To calculate the info measure we used the ratio of observed variance of the dosages to the expected binomial variance at HardyWeinberg equilibrium, as used in the imputation software MACH [Scott et al., 2007]. Supplementary Figure 3a, 3b and 4a, 4b show that in scenario 5, with a quantitative trait, there is also increased statistical power and less bias when using the full genotype probabilities. It also show that when keeping individuals with 0 reads, there is less bias, but the same statistical power. In Table 3 we run the analysis from Table 2, but including individuals with no reads. In this scenario the difference between using dosages and genotype probabilities has been almost erased. However it is worth noticing that in this scenario, expect for the true genotype, we lose statistical power compared to when we remove individuals without reads. To further investigate these scenarios we looked at the bias of the estimated effect sizes, as Supplementary Figure 9 and 10 show for a RR of 1.14 we see that the EM model provides less biased estimates of the effect sizes compared to dosages, for both scenarios.

**Table 2:** The statistical power for different simulated effect sizes or relative risk (RR) of the genotype. The phenotype is simulated as a binary trait. We have done 10,000 simulations for each tested effect size. The casual allele of the genetic variant has a frequency of 0.05 and the disease has a prevalence of 0.10 in the population, we use a significance threshold of 10^*–*5^. We have used the sample allele frequency prior as there is no population structure. We have 50,000 controls and cases with an average sequencing depth of 1*X* and 4*X* respectively, here we have removed individuals with 0 reads (0*X*). The *R*^2^ values denote how well the true genotype is predicted with this data, they are calculated like the info measure used in the MACH imputation software [Scott et al., 2007]

| data / RR: | 1 | 1.1 | 1.12 | 1.14 | 1.16 |
| --- | --- | --- | --- | --- | --- |
| True genotype | 0 | 0.587 | 0.868 | 0.978 | 0.999 |
| Dosage | 0 | 0.114 | 0.300 | 0.431 | 0.808 |
| Genotype probs. | 0 | 0.163 | 0.388 | 0.659 | 0.862 |
| $R^2$ cases/controls | 0.91/0.85 | 0.91/0.85 | 0.91/0.84 | 0.90/0.84 | 0.90/0.84 |

**Table 3:** This Table is similar to Table 2 it is the same scenario also, but where we include individuals with 0 reads.

| data / RR: | 1 | 1.1 | 1.12 | 1.14 | 1.16 |
| --- | --- | --- | --- | --- | --- |
| True genotype | 0 | 0.813 | 0.974 | 0.999 | 1.000 |
| Dosage | 0 | 0.0914 | 0.262 | 0.523 | 0.772 |
| Genotype probs. | 0 | 0.0974 | 0.273 | 0.538 | 0.783 |
| $R^2$ cases/controls | 0.91/0.77 | 0.90/0.75 | 0.90/0.75 | 0.90/0.75 | 0.90/0.75 |

Supplementary Figure 5a, 5b and 6a, 6b, show that in scenario 6, with a quantitative trait there is increased statistical power and less bias of, when using genotype probabilities compared to dosages. Lastly if we do scenario 6, but for a binary phenotype we show increased statistical power and a smaller bias of the effect size, when using genotype probabilities compared to dosages as shown in Supplementary Figure 7a, 7b and 8a, 8b. And that removing individuals with 0 reads, leads to increased statistical power and less bias.

### 4.3 Comparison with SNPTEST

SNPTEST [Marchini et al., 2007] also implements an EM algorithm for doing association (using the *-method em*) with genotype probabilities also using a generalised linear model framework. To compare the performance of the two methods the bias of the estimated effect sizes and statistical power were investigated, Figure 4 shows that SNPTEST’s effect sizes are downward biased, whereas `ANGSD-asso`’s EM model has no bias, when using an individual allele frequency prior, and increased statistical power compared to SNPTEST. We found that this only happens in scenarios with adjustment for confounders, like this scenario where covariates are included to adjust for population structure. This happens as SNPTEST adjusts for covariates by first regressing the covariates (plus an intercept) on the phenotype, and then subsequently running the EM algorithm with intercept and genotype in the linear model on the residuals. SNPTEST performs worse when using an individual allele frequency prior, since this is more correlated with ancestry than when using a sample frequency prior. Correlation between variables in a linear model, and then using the technique of regressing out covariates leads to biased parameter estimates [Freckleton, 2002]. When not including covariates the estimates of the effect sizes are the same (data not shown). When using the SNPTEST approach for dosages (using the *-method expected*) the effect sizes are the same, also when including covariates (data not shown). We have used the most recent version of SNPTEST (v2.5.4-beta3).

**Figure 4:**
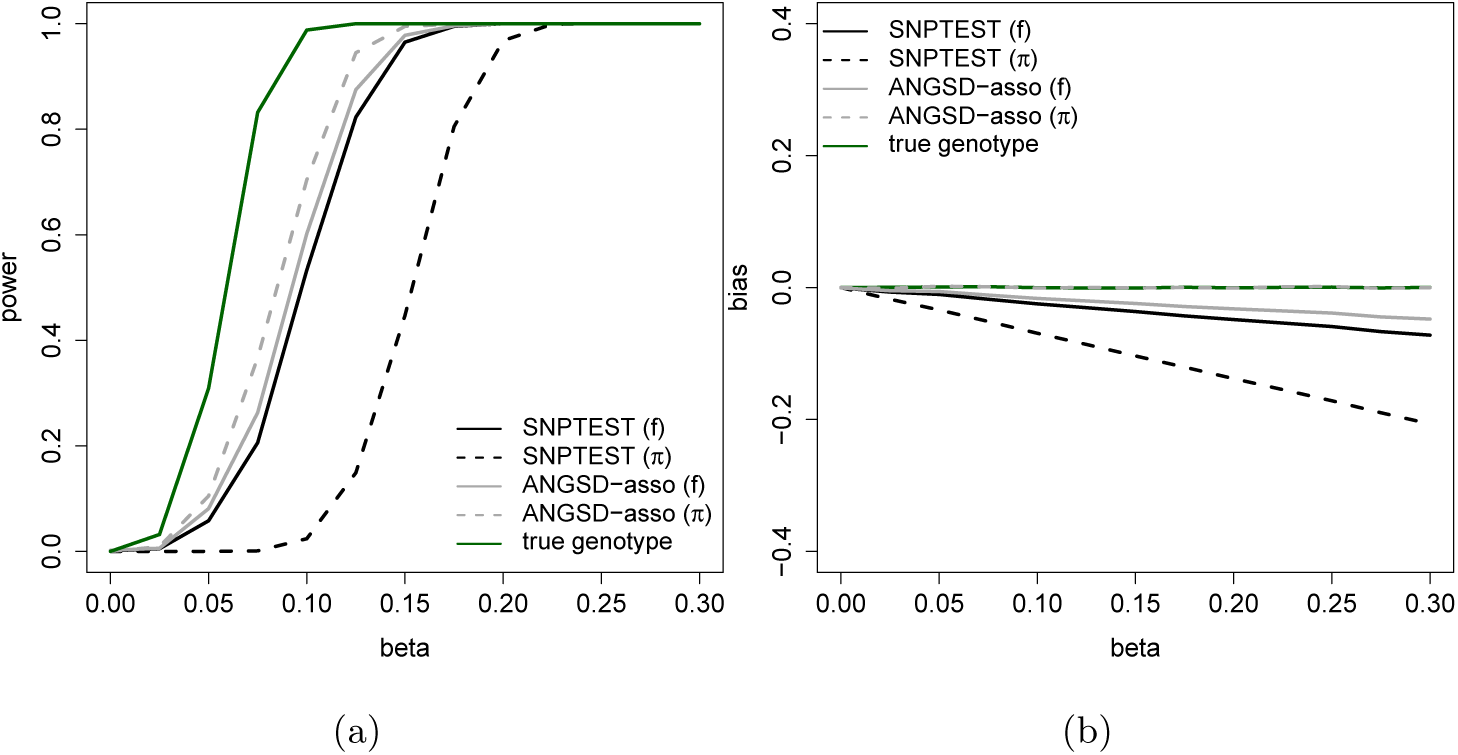
This data is simulated with 10,000 individuals with an average depth of 0.1, 1, 10, 20*X*, 2,500 individuals each. Varying the effect size of the genotype (*β*). There is an effect of ancestry of population 1 (*γ* = 0.5), the admixture proportions for population 1 are for the same 2,500 individuals each (*Q*_1_=0.15, 0.4, 0.95, 1). We use a significance threshold of 10^*–*3^. The linear models are adjusted for ancestry, SNPTEST was run without transforming the phenotype or covariates to make it as comparable to `ANGSD-asso`’s EM model as possible. Each point is based on 1,000 simulations. **(a)**: We show the statistical power to detect a true association using `ANGSD-asso`’s EM model and SNPTEST’s EM model respectively with a sample frequency prior (f) and an individual allele frequency prior (*π*). **(b)**: We show the bias of our estimated effect size.

We also compared `ANGSD-asso`’s EM model to SNPTEST in terms of computational speed and found that our EM algorithm converges faster. Also it can be run multi-threaded resulting in much reduced run times.

`ANGSD-asso`’s EM model is faster than SNPTEST, especially for binary data, as shown in Figure 5 and for quantitative data as shown in Supplementary Figure 12. `ANGSD-asso`’s EM model is capable of analysing data sets of 100,000 individuals in less than 10 hours. Our hybrid approach can handle the analyses in less than 17 hours unthreaded. SNPTEST will takes days to run the largest data set, when running a logistic model.

**Figure 5:**
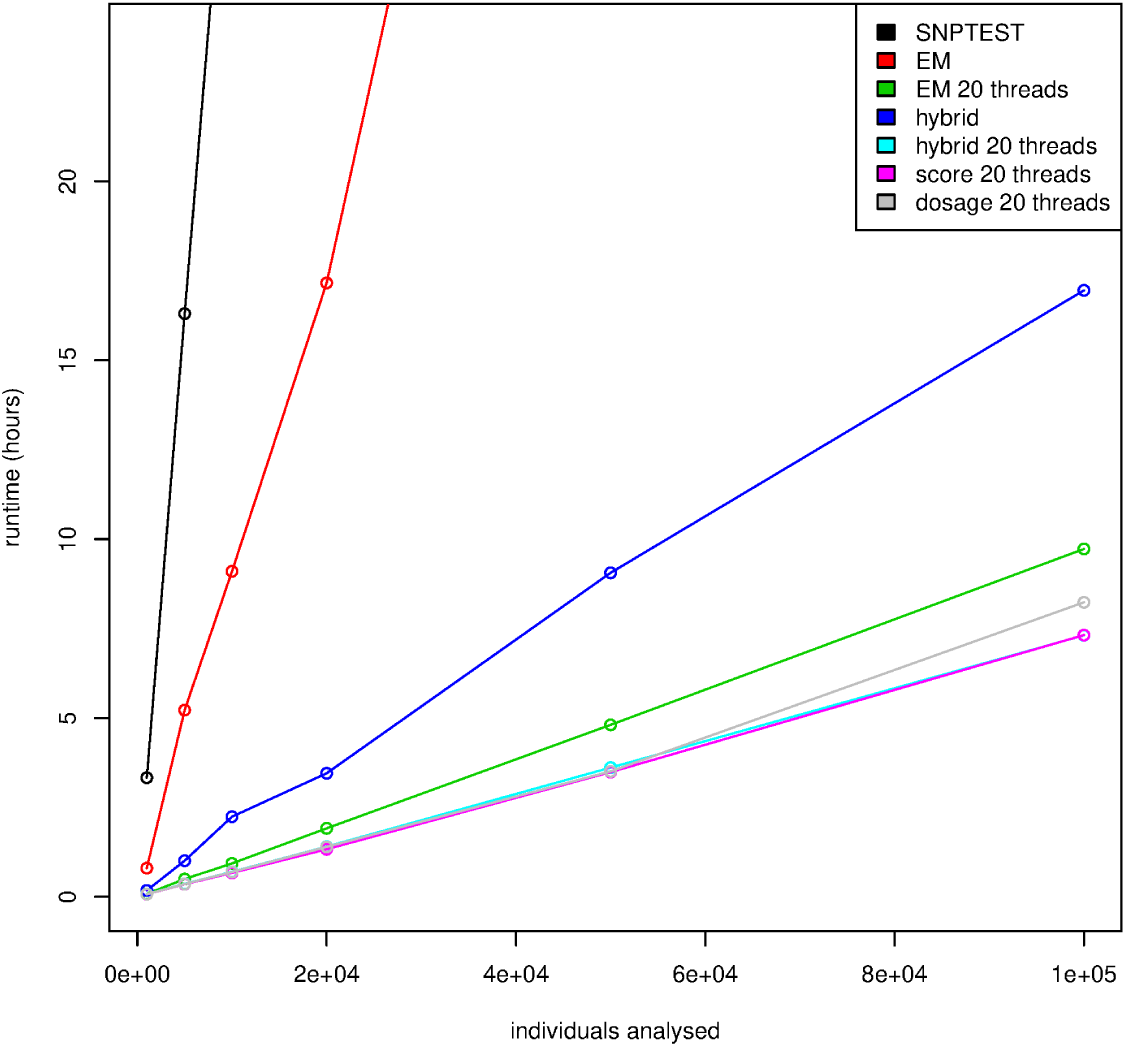
Running times for an analysis of a simulated binary trait with 442,769 genetic variants, varying the number of individuals (1,000, 5,000, 10,000, 20,000, 50,000 and 100,000), the model is run with 2 covariates (age and gender). The genetic data has an average depth of 1*X*. For each point we have run the analysis 3 times and then used the mean running time. All runs but the SNPTEST analysis is run in `ANGSD`.

In order to achieve faster convergence of `ANGSD-asso`’s EM model, we first do regression on the genotype dosages. We then use the coefficients obtained from the dosage regression as the starting guess for the coefficients for the EM algorithm (we refer to this as priming). As shown in Supplementary Figure 11 this drastically reduces the number of iterations needed for convergence of the EM algorithm.

## 5 Discussion

### 5.1 Implementation of model

We have implemented `ANGSD-asso`’s EM model for taking genotype uncertainty into account when doing association studies. The advantage of this approach compared to the score test [Skotte et al., 2012], is that the effect size of the unobserved genotype is estimated. The effect size helps provide further insights into the relationship between genotype and phenotype. Furthermore the estimated effect sizes also means we can make use of LD-score regression [Bulik-Sullivan et al., 2015]. It is shown through simulations that `ANGSD-asso`’s EM model has increased statistical power and less bias compared to SNPTEST as shown in Figure 4, when including covariates in the model, due to SNPTEST adjusting for covariates by first regressing them on the phenotype and then running the EM algorithm on the residuals. Including covariates in the linear model is a common way to deal with confounders in association studies.

### 5.2 Different priors in structured and homogeneous populations

We have shown how using an individual allele frequency prior, when estimating genotype probabilities, gives better statistical power to detect an association, when dealing with NGS data with population structure as shown in Figure 3. Also it removes issues with an increased false positives rate when there is sequencing depth phenotype correlation as shown in Figure 2. This correlation might arise if the sequencing is not randomised, for example if cases and controls are being sequenced at different places thereby creating a systematic bias, or if different cohorts have been sequenced at different places. The scenarios from Table 1 are most likely to arise when dealing with non model species where imputation cannot be done. This leads us to recommend using an individual allele frequency prior when doing association studies with NGS data in structured populations, where imputation is not possible.

### 5.3 Comparison with dosages in large scale studies

In Table 2 and 3 we show through simulations increased statistical power when using genotype probabilities compared to dosages, with a larger gain in power for the scenario from Table 2. In both instances it is a case control study with low depth sequencing data, where cases and controls have different average sequencing depths. A scenario like this, where there is better genotypic information for some individuals, could arise doing imputation. As shown in Table 2 and 3 with the info measure (*R*^2^) for controls and cases, where cases have more informative genetic data. This could happen if a certain population is not being represented in the reference panel used for imputation or if different reference panels or SNP-chips are used for cases and controls. A systematic difference in imputation quality is roughly equivalent to having a different average sequencing depth. We also show increased statistical power and a smaller bias of the effect size, when using genotype probabilities compared to dosages, when analysing a quantitative trait, as shown in Supplementary Figure 3a, 3b and 4a, 4b (scenario 5 from Table 1) even though this is a much smaller study in terms of the number of individuals. Furthermore in Supplementary Figure 5a, 5b and 6a, 6b (scenario 6 from Table 1), we show increased statistical power and a smaller bias of the effect size, when using genotype probabilities compared to dosages, when there is also population structure. Lastly for scenario 6 from Table 1, but for a binary phenotype we show increased statistical power and a smaller bias of the effect size, when using genotype probabilities compared to dosages as shown in Supplementary Figure 7a, 7b and 8a, 8b.

Another conclusion we can draw is that when there is sequencing depth and phenotype correlation, our estimates of the effect size will be biased when we do not know the true frequency as shown in Supplementary Figure 4, 6 and 8. Furthermore when there is sequencing depth and phenotype correlation and when analysing a quantitative trait, we should keep individuals with 0 reads as that leads to a smaller bias, as shown in Supplementary Figure 4 and 6, even when using true genotypes. For a binary phenotype we should remove individuals with 0 reads, as that leads to a smaller bias as shown in Supplementary Figure 8, 9 and 10.

In this article we have not explored how imputation might affect association in a structured population. With `ANGSD-asso` implemented in ANGSD we have made it possible to do large association studies with low depth sequencing data retaining maximal statistical power, and also estimating effect sizes. SNPTEST is too slow for the analysis of large scale data sets. The speed-up of `ANGSD-asso`’s EM model compared to SNPTEST is due to priming for faster convergence of the EM algorithm and threaded analysis using the ANGSD [Korneliussen et al., 2014] framework. `ANGSD-asso` makes the analysis of large scale data possible as done in Liu et al. [2018] (141,431 individuals) while retaining maximal statistical power.

