## Supplementary Material for "Efficient approaches for large scale GWAS studies with genotype uncertainty"

### 1 Simulating scenario 1 and 2

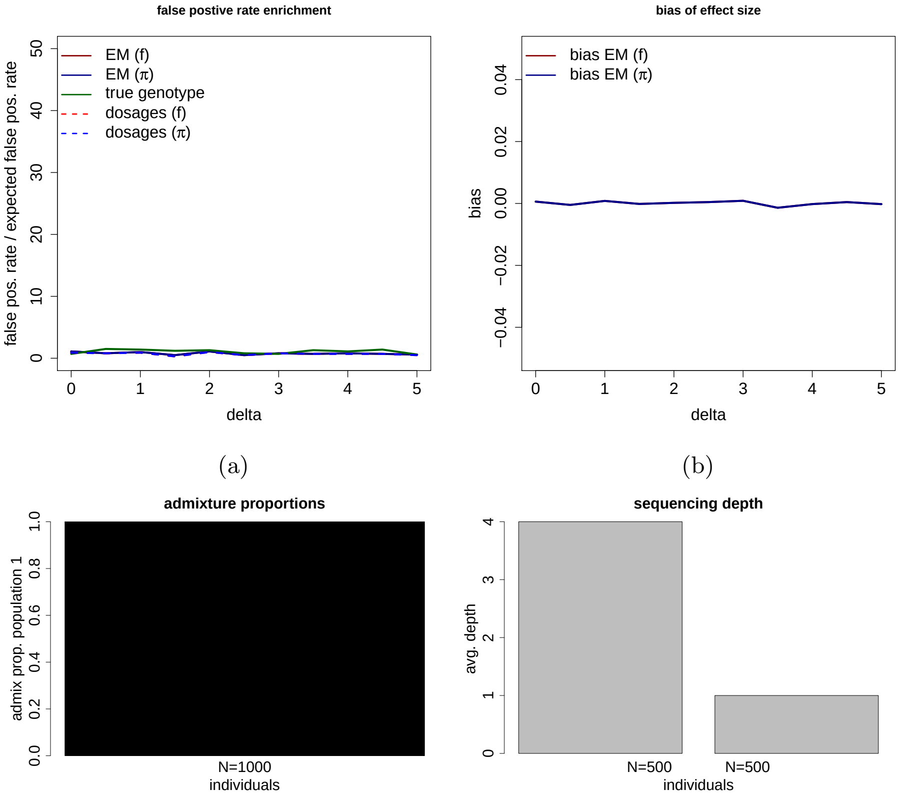


(c)

Figure 1: This data is simulated according to scenario 1 in Table 1 varying the sequencing depth and phenotype correlation ($\delta$) (eq. 12 in the main text). We have a population with 1,000 individuals without structure. We use a significance threshold of $10^{-3}$. Each point is the mean value from 10,000 simulations. **(a)**: We show the false positive rate divided by the expected false positive rate ($10^{-3}$) as a function of the sequencing depth phenotype correlation, using `ANGSD-asso`'s EM model and dosage model respectively with a sample frequency prior (f) and an individual allele frequency prior ($\pi$). **(b)** We show the bias of our estimated effect size. **(c)** The simulated admixture proportions and the mean sequencing depth for the simulated individuals.

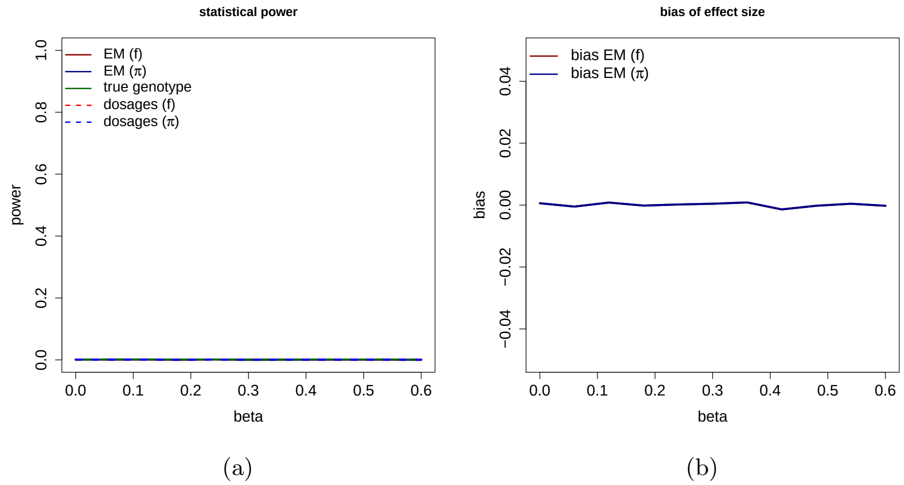


Figure 2: This data is simulated according to scenario 2 in Table 1 varying the effect size of the genotype ($\beta$). We have the same mean sequencing depth as in Supplementary Figure 1 (c). The phenotype is simulated as a quantitative trait, with different effect sizes of the genotype, for each tested effect size it is the mean value from 10,000 simulations. **(a)**: We show the statistical power to detect a true association with a significance threshold of $(10^{-3})$, using `ANGSD-asso`'s EM model and dosage model respectively with a sample frequency prior (f) and an individual allele frequency prior ($\pi$). **(b)**: We show the bias of our estimated effect size.

#### 2 Sequencing depth phenotype correlation with a quantitative phenotype

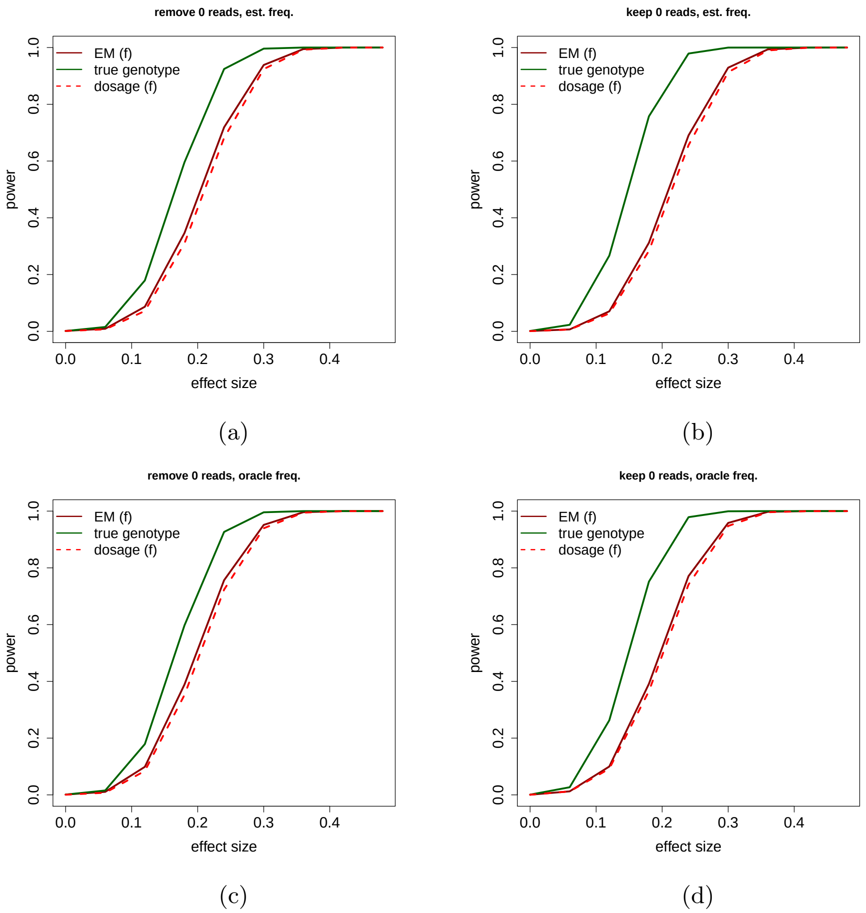


Figure 3: This data is simulated according to scenario 5 in Table 1 varying the effect size of the genotype ($\beta$), sequencing depth phenotype correlation ($\delta$) is fixed at a value of 5 (eq. 12 in the main text). The phenotype is simulated as a quantitative trait, with different effect sizes of the genotype, for each tested effect size it is the mean power from 10,000 simulations. **(a)**: We show the statistical power to detect a true association with a significance threshold of ($10^{-3}$), using `ANGSD-asso`'s EM model and dosage model respectively with a sample frequency prior (f), removing individuals with 0 reads and estimating $f$ from the genotype likelihoods. **(b)**: Like (a) but keeping individuals with 0 reads. **(c)**: Like (a) but knowing the simulated $f$. **(d)**: Like (a) but keeping individuals with 0 reads and knowing the simulated $f$.

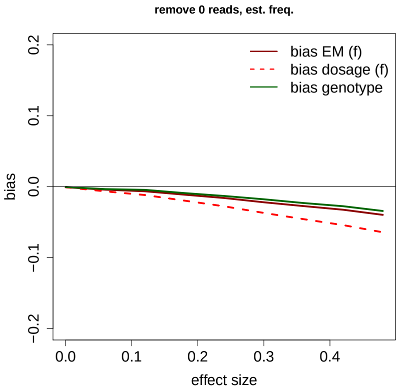


(a)

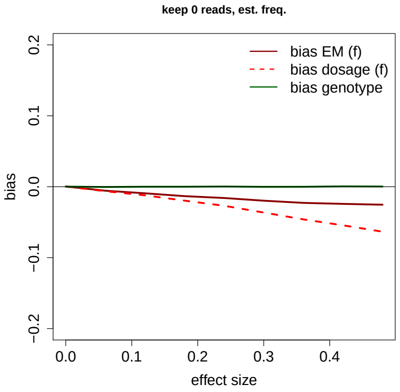


(b)

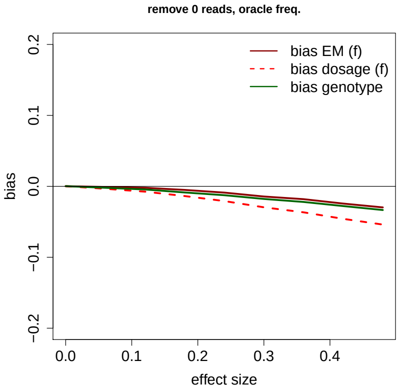


(c)

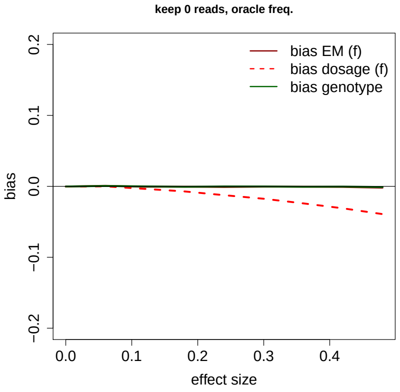


(d)

Figure 4: Like Supplementary Figure 3, but showing the bias of the estimated effect size.

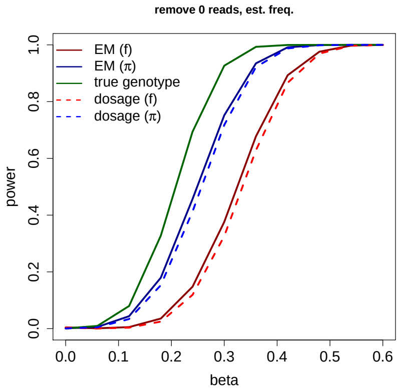


(a)

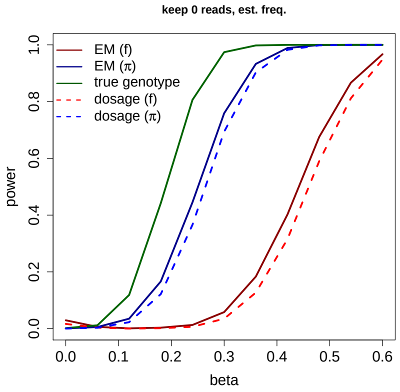


(b)

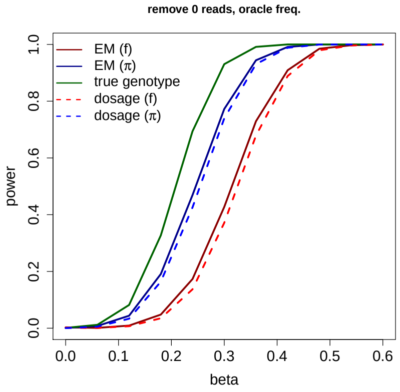


(c)

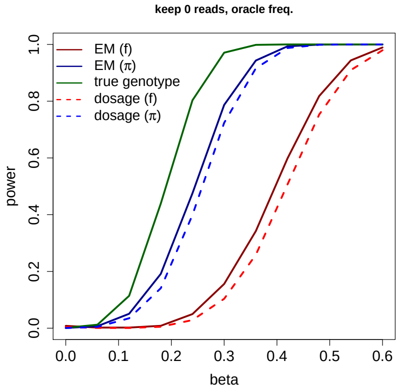


(d)

Figure 5: This data is simulated according to scenario 6 in Table 1 varying the effect size of the genotype ($\beta$), sequencing depth phenotype correlation ($\delta$) is fixed at a value of 5 (eq. 12 in the main text). There is an effect of ancestry of population 1 ($\gamma = 1$). The phenotype is simulated as a quantitative trait, for each tested effect size it is the mean power from 10,000 simulations. **(a)**: We show the statistical power to detect a true association with a significance threshold of ($10^{-3}$), using `ANGSD-asso`'s EM model and dosage model respectively with a sample frequency prior (f) and an individual allele frequency prior ($\pi$). **(b)**: Like (a) but keeping individuals with 0 reads. **(c)**: Like (a) but knowing the simulated $f$ and $\pi$. **(d)**: Like (a) but keeping individuals with 0 reads and knowing the simulated $f$ and $\pi$.

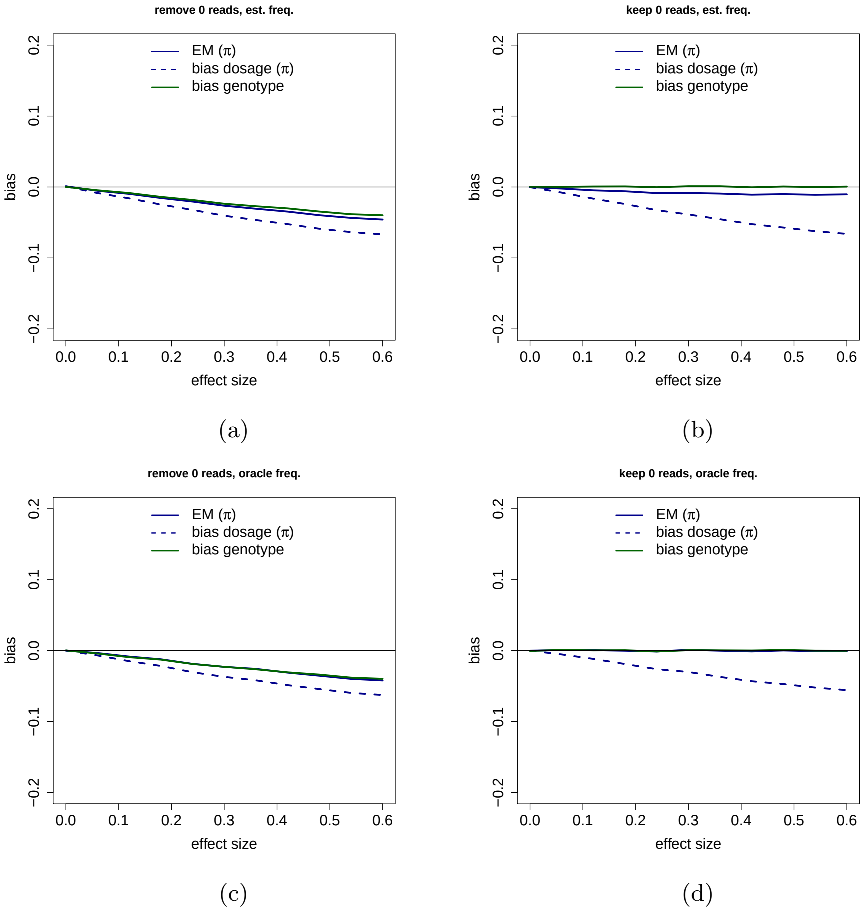


Figure 6: Like Supplementary Figure 5, but showing the bias of the estimated effect size.

### 3 Sequencing depth phenotype correlation with a binary phenotype and population structure

The binary phenotype is simulated like this:

$$p(y = 1) = \frac{1}{1 + e^{-(\beta \cdot g + \gamma \cdot q)}} \tag{1}$$

Depth phenotype correlation was simulated like this:

$$p(D_{high}|y = 1) = \frac{1}{1 + e^{-\delta}} \tag{2}$$

$$p(D_{high}|y = 0) = 1 - \frac{1}{1 + e^{-\delta}} \tag{3}$$

The higher $\delta$ is the more probable cases will be in the high depth category.

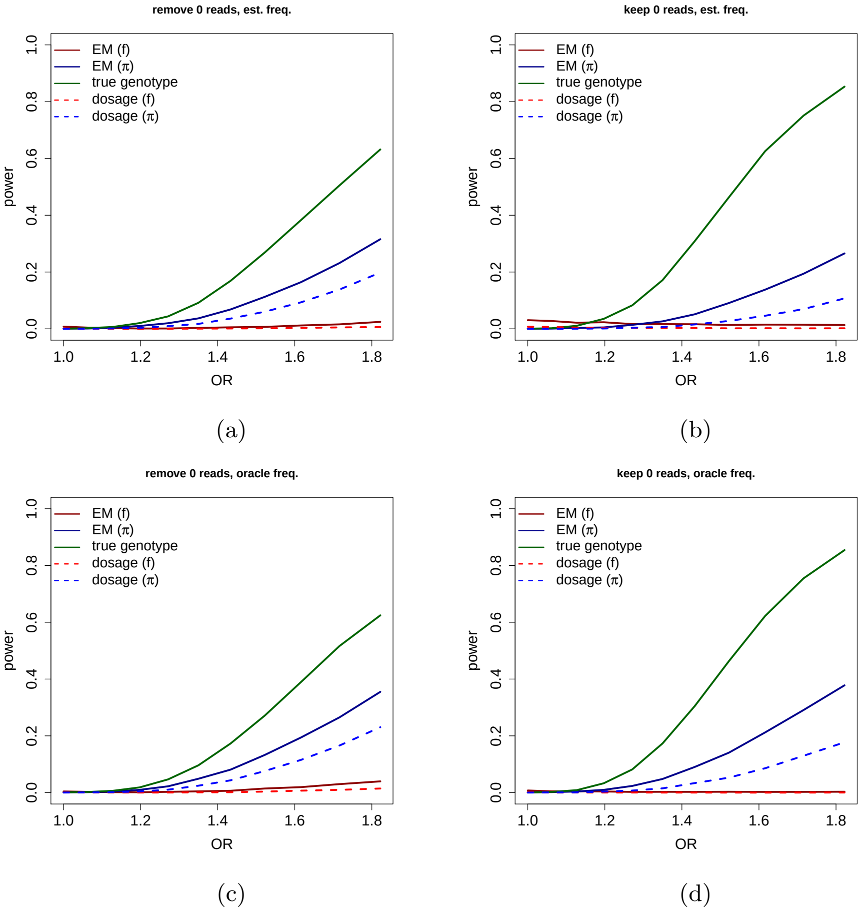


Figure 7: This data is simulated according to scenario 6 in Table 1 varying the effect size of the genotype ($\beta$), sequencing depth phenotype correlation ($\delta$) is fixed at a value of 5 (eq. 3). There is an effect of ancestry of population 1 ($\gamma = 1$). The phenotype is simulated as a binary trait according to eq. 1, with different effect sizes of the genotype, for each tested effect size it is the mean power from 10,000 simulations. **(a)**: We show the statistical power to detect a true association, with a significance threshold of ($10^{-3}$) using `ANGSD-asso`'s EM model and dosage model respectively with a sample frequency prior (f) and an individual allele frequency prior ($\pi$). **(b)**: Like (a) but keeping individuals with 0 reads. **(c)**: Like (a) but knowing the simulated $f$ and $\pi$. **(d)**: Like (a) but keeping individuals with 0 reads and knowing the simulated $f$ and $\pi$.

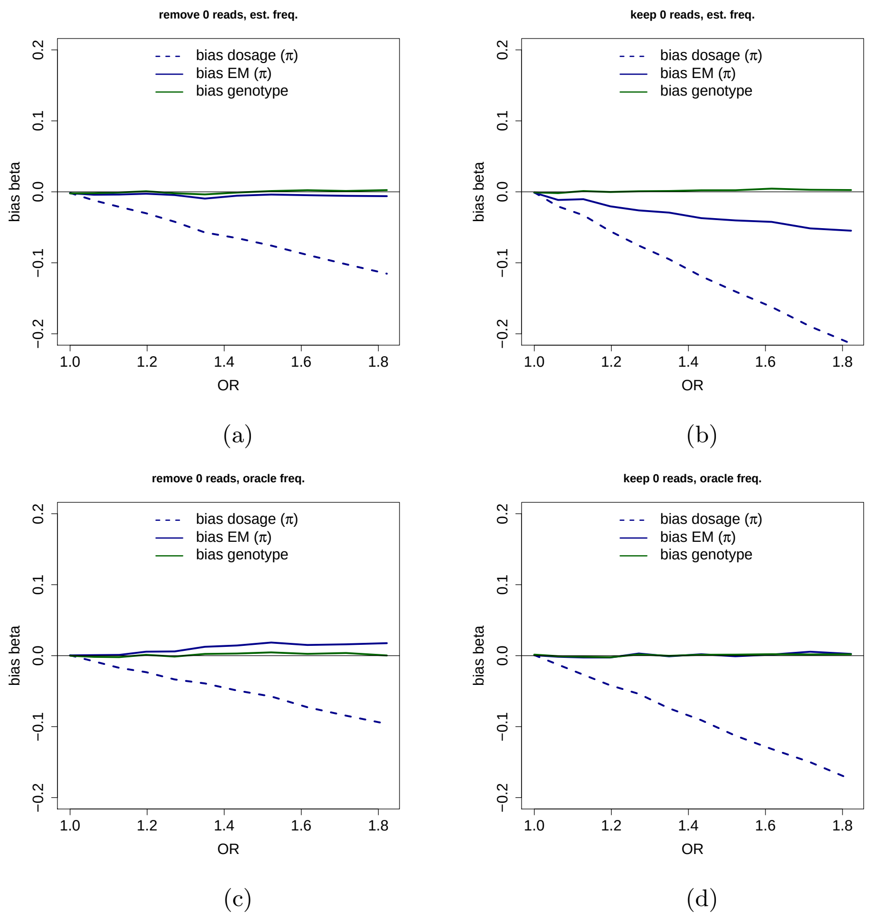


Figure 8: Like Supplementary Figure 7, showing the bias of the estimated effect size ($\beta$) not the odds ratio.

### 4 Sequencing depth phenotype correlation with a binary phenotype

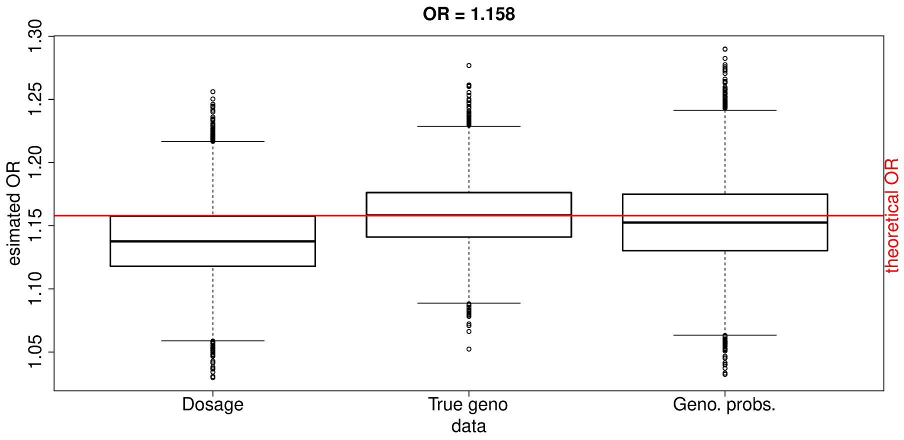


Figure 9: Estimated odds ratio (OR) from all 3 methods used in Table 2, in order to show the bias of the estimated effect size. The estimated effect sizes were simulated using a relative risk (RR) of 1.14. For showing the bias of the estimated effect size, the OR is used, as this can be obtained from the logistic regression. A RR of 1.14 with a disease prevalence of 0.1 is equivalent to an OR of 1.158. The formula for converting OR to RR from Zhang and Kai [1998] was used.

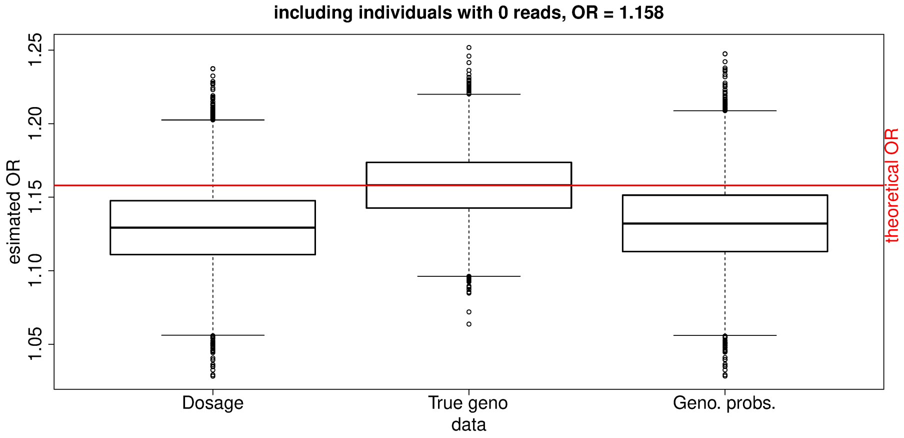


Figure 10: Like Supplementary Figure 9, but keeping individuals with 0 reads, like in Table 3.

#### 4.1 Priming coefficients for faster convergence

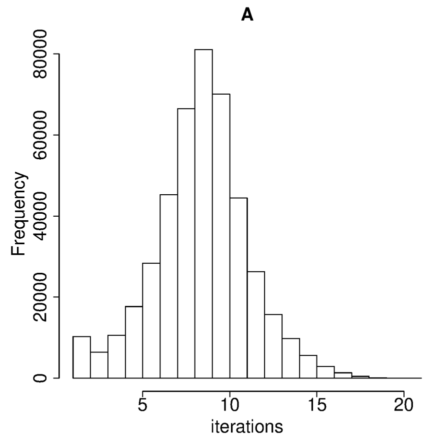


(a)

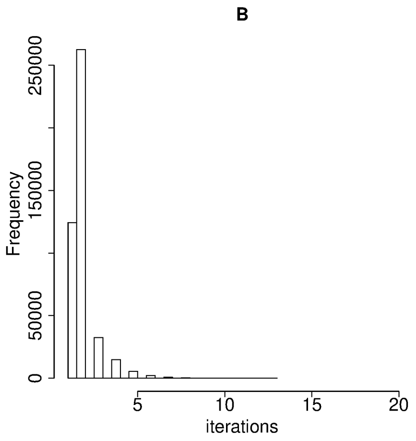


(b)

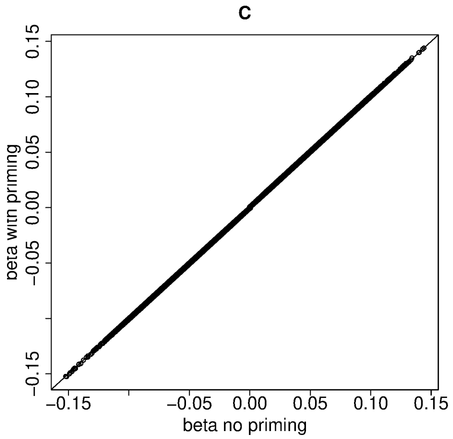


(c)

Figure 11: **(a)**: Iterations for the analysis of 442,769 sites for 5,000 individuals with a simulated quantitative trait. Sequencing depth is on average $1X$ for the individuals when not priming coefficients. **(b)**: Same analysis as **(a)** but when priming coefficients. **(c)**: Effect sizes for `ANGSD-asso`'s EM model when not using priming and when using priming.

#### 4.2 Genotype likelihoods from NGS data

Next generation sequencing (NGS) produces reads with the observed nucleotide bases. These reads are then aligned to a reference genome, and each position will be covered by a certain number of reads, the more reads covering a position the more certainty there is of the true genotype of that position. The number of reads covering a position is called the sequencing depth. Each base in a sequenced read will have a quality score denoting the certainty of the called nucleotide base. Genotype likelihoods (GLs) is the likelihood of observing the sequence data $x_j$ given an unknown genotype $G$, meaning $p(x_j|G)$ for a given position in the genome $j$. The sequence data $x_j$ will consist of $R$ observed bases $x_j = (b_1, b_2, ..., b_R)$.

The GL $p(x_j|G)$ is calculated using the quality scores of $x_j$, one method for doing so is by assuming the $R$ observed bases are independent given our genotype $G$

$$p(X_j|G) \propto \prod_{r=1}^{R} p(b_r|G) = \prod_{r=1}^{R} p(b_r|A_1, A_2) =$$

Where $A_1$ and $A_2$ are the two alleles of the genotype $G$, summing over the alleles $A$ and rewriting the likelihood

$$\prod_{r=1}^{R} \sum_{A \in A_1, A_2} p(b_r|A)p(A) =$$

And since $p(A)$ or the probability of observing one of the two alleles must be one half.

$$\prod_{r=1}^{R} \left( \frac{1}{2} p(b_r|A_1) + \frac{1}{2} p(b_r|A_2) \right).$$

The probability of a base $b$ given an allele $A$ relates to the error rate $\epsilon$.

$$p(b|B) = \begin{cases} \frac{\epsilon}{3}, & b \neq A \\ 1 - \epsilon, & b = A \end{cases}$$

Here $R$ is the depth at site $j$, $b_r$ is the observed $r$th base, and $\epsilon$ is the probability of an error as calculated from the quality score of $b_r$.

This is the approach used in the GATK framework McKenna et al. [2010], other approaches where the quality score is not used directly exist, for example SAMtools Li et al. [2009].

The likelihood can be used to calculate the posterior probability of the genotype $G$ or the genotype probability, using a genotype prior, $p(G)$ and Bayes Theorem, like in eq. 8 in the main text.

### 5 Poisson distribution written as an exponential family

For the Poisson distribution we have that the probability of our phenotype $y_i$ of the $i$th individuals with value $k$ can be written like this

$$p(y_i = k|G_i, \mathbf{z_i}) = \frac{\lambda_i^k e^{-\lambda_i}}{k!}. \tag{4}$$

Where our linear predictor for the Poisson regression is

$$\eta_i = \log E[y_i] = \log \lambda_i. \tag{5}$$

We can write up $\exp(\log(p(y_i = k|G_i, \mathbf{z_i}))$

$$\exp\left(\log\left(\frac{\lambda_i^{y_i} e^{-\lambda_i}}{y_i!}\right)\right) = \exp(\log \lambda_i^{y_i} e^{-\lambda_i} - \log(y_i!)) = \tag{6}$$

$$\exp(y_i \log \lambda_i + \log e^{-\lambda_i} - \log(y_i!)) = \exp(y_i \log \lambda_i - \lambda_i - \log(y_i!)). \tag{7}$$

Then we can write it as an exponential family using notation from Dobson and Barnett [2008]

$$\exp\left(\frac{y_i\eta_i - b(\eta_i)}{a(\phi)} + c(y_i, \phi)\right). \tag{8}$$

Giving us $a(\phi) = 1, b(\eta_i) = \lambda_i = \exp(\eta_i)$ and $c(y_i, \phi) = -\log(y!)$.

### 6 EM algorithm in ANGSD-asso's EM model

Starting from eq. 4 in the main text where we have

$$\sum_i^N \log\left(\sum_{g\in\{0,1,2\}} p(y_i|G_i = g, \mathbf{z_i}, \theta)p(G_i = g|x_i)\right). \tag{9}$$

The term $p(y_i|G_i, \mathbf{z_i}, \theta)$ we base on a linear regression model. We assume that given the genotype $G$, covariates $Z$ and parameters $\theta$ the phenotype follow a normal distribution with a mean given by

$$\eta_i = \alpha + \beta G_i + \sum_c \gamma_c z_{ic}. \tag{10}$$

We can then apply the EM algorithm for maximising the likelihood. First we do the E-step, the term $p(y_i|G_i, z_i, \theta)$ is the only one that depends on $\theta$, this is equivalent to maxisming

$$E_{G|\mathbf{y},\mathbf{x},Z,\theta}[\log p(\mathbf{y}|G_i,Z,\theta)] = E_{G|\mathbf{y},\mathbf{x},Z,\theta}[\log \prod_i^N p(y_i|G_i,\mathbf{z_i},\theta)] \tag{11}$$

$$= \sum_i^N E_{G|y_i,x_i,\mathbf{z_i},\theta}[\log p(y_i|G_i,\mathbf{z_i},\theta)]. \tag{12}$$

And then the M-step is differentiating this as in Lake et al. [2003]

$$\frac{\partial}{\partial \beta}\sum_i^N E_{G|y_i,x_i,\mathbf{z_i},\theta}[\log p(y_i|G_i,\mathbf{z_i},\theta)] \tag{13}$$

$$= \sum_i^N E_{G|y_i,x_i,\mathbf{z_i},\theta}\left[\frac{\partial}{\partial \beta}\log p(y_i|G_i,\mathbf{z_i},\theta)\right] \tag{14}$$

$$= \sum_i^N E_{G|y_i,x_i,\mathbf{z_i},\theta}\left[\frac{\partial \eta_i}{\partial \beta}\frac{\partial}{\partial \eta_i}\log p(y_i|G_i,\mathbf{z_i},\theta)\right] \tag{15}$$

Using the rule of differentiating a sum going from (13) to (14). Using the chain rule going from (14) to (15). ($\frac{dz}{dx} = \frac{dz}{dy}\frac{dy}{dx}$)

The term $p(y_i|G_i,\mathbf{z_i},\theta)$ can be written as an exponential family

$$p(y_i|G_i,\mathbf{z_i},\theta) = \frac{1}{\sqrt{2\pi\sigma^2}}\exp\frac{-(y-\eta_i)^2}{2\sigma^2}$$

$$= \exp\left(\frac{y_i\eta_i - \eta_i^2/2}{\sigma^2} - y_i^2/2\sigma^2 - \log(2\pi\sigma^2)/2\right) = \exp\left(\frac{y_i\eta_i - b(\eta_i)}{a(\sigma)} + c(y_i,\sigma)\right).$$

Where $a(\sigma) = \sigma^2$, $b(\eta_i) = \eta_i^2/2$ and $c(y_i,\sigma) = -y_i^2/2\sigma^2 - \log(2\pi\sigma^2)/2$. For a logistic regression we have $a(\sigma) = 1$, $b(\eta_i) = \log(1+e^{\eta_i})$ and $c(y_i,\sigma) = 0$. And for a Poisson regression we have $a(\phi) = 1, b(\eta_i) = \lambda_i = \exp(\eta_i)$ and $c(y_i,\phi) = -\log(y!)$.

In the case of $b(\eta_i) = \eta_i^2/2$ and $a(\sigma) = \sigma^2$ (where $b'(\eta_i) = \eta_i$), we can derive the following.

$$\sum_i^N E_{G|y_i,x_i,\mathbf{z_i},\theta}\left[\frac{\partial\eta_i}{\partial\beta}\frac{\partial}{\partial\eta_i}\frac{y_i\eta_i - b(\eta_i)}{a(\sigma)} + c(y_i,\sigma)\right] \tag{16}$$

$$= \sum_i^N E_{G|y_i,x_i,\mathbf{z_i},\theta}\left[\frac{\partial\eta_i}{\partial\beta}\frac{y_i - b'(\eta_i)}{a(\sigma)}\right] \tag{17}$$

$$= \sum_i^N E_{G|y_i,x_i,\mathbf{z_i},\theta}\left[G_i\frac{y_i - \eta_i}{\sigma^2}\right] \tag{18}$$

$$= \sum_i^N \sum_{g\in\{0,1,2\}}\left(G_i\frac{y_i - \eta_i}{\sigma^2}\right)p(G_i = g|y_i, x_i, \mathbf{z_i}, \theta). \tag{19}$$

In going from (18) to (19) we are taking the expectation across all values of $G$. For the other variables ($\alpha$, $\gamma$) in (10)

$$\sum_i^N \sum_{g\in\{0,1,2\}}\left(z_i\frac{y_i - \eta_i}{\sigma^2}\right)p(G_i = g|y_i, x_i, \mathbf{z_i}, \theta). \tag{20}$$

Where for $\alpha$ we will just replace $z_i$ with 1.

We recognise (19) and (20) as the score functions of a weighted regression, with regards to the respective terms ($\alpha$, $\beta, \gamma$) [Dutang, 2017]. Each individual $i$, contributes one observation per possible genotype $G$ the weights are given by $p(G_i|y_i, x_i, \mathbf{z_i}, \theta)$ which is the probability of a genotype $G$ given the phenotype $y$, covariates $\mathbf{z}$ and parameters $\theta$. This is maximised by doing weighted least squares, where the parameters $\theta$ are chosen to maximise the likelihood.

The term $p(G_i|y_i, x_i, \mathbf{z_i}, \theta)$ can be estimated using Bayes' theorem again making use of the assumption $p(G_i|x_i, \mathbf{z_i}, \theta) = p(G_i|x_i)$, and that we can ignore the sequence data when we have the genotype $p(y_i|G_i, x_i, \mathbf{z_i}, \theta) = p(y_i|G_i, \mathbf{z_i}, \theta)$ this yields

$$p(G_i|y_i, x_i, \mathbf{z_i}, \theta) = \frac{p(y_i|G_i, \mathbf{z_i}, \theta)p(G_i|x_i)}{\sum_{g\in\{0,1,2\}} p(y_i|G_i = g, \mathbf{z_i}, \theta)p(G_i = g|x_i)}. \tag{21}$$

#### 6.1 Optimisation strategy for a normal distributed phenotype

First an initial guess of the standard deviation is calculated from the phenotype, using the sample standard deviation

$$s = sd[y] \tag{22}$$

Then linear regression is done with the full model, using the dosages calculated form the genotype probabilities, to estimate the an initial guess of the coefficients for `ANGSD-asso`'s EM model for faster convergence. This is referred to in this article as priming the coefficients.

Regression weights are then calculated according to (21) and weighted least squares is done using the parameters and weights from the last iteration of the EM algorithm. Each individual has three entries in the design matrix $(\mathbf{G}, Z)$, where $\mathbf{G}$ is a vector with each of the three possible genotypes for each individual, each weighted by $p(G_i|y_i, x_i, \mathbf{z_i}, \theta)$ as estimated for that individual and for that genotype. Then $s$ is updated by using the weighted sum of squared residuals, from the weighted least squares with $n - o$ degrees of freedom. Where $n$ is the number of individuals and $o$ is the number of coefficients in the linear model as described in (10). The term $p(y_i|G_i, \mathbf{z_i}, \theta)$ can be calculated using a normal distribution with the following parameters, where $\eta_i$ is from (10).

$$p(y_i|G_i, \mathbf{z_i}, \theta) = \mathcal{N}(\eta_i, s^2). \quad (23)$$

### 7 Running times for a quantitative trait

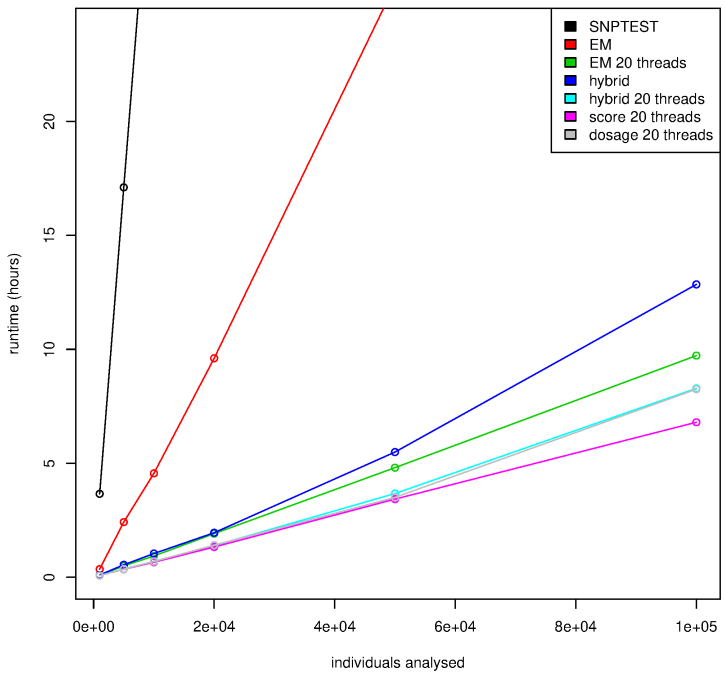


Figure 12: Running times for an analysis of a simulated quantitative trait with 442,769 genetic variants, varying the number of individuals (1,000, 5,000, 10,000, 20,000, 50,000 and 100,000), the model is run with 2 covariates (age and gender). The genetic data has an average depth of $1X$. For each point we have run the analysis 3 times and then used the mean running time. All runs but the SNPTEST analysis is run in `ANGSD`.
